# Enteric glial cells respond to a dietary change in the lamina propria in a MyD88-dependent manner

**DOI:** 10.1101/2020.11.23.394601

**Authors:** Zhuanzhuan Liu, Hongxiang Sun, Ming Liang, Jing Gao, Liyuan Meng, Xingping Zheng, Yanxia Wei, Yanbo Kou, Yugang Wang

**Affiliations:** Laboratory of Infection and Immunity, Jiangsu Key Laboratory of Immunity and Metabolism, Department of Pathogenic Biology and Immunology, Xuzhou Medical, University, Xuzhou, Jiangsu Province, China 221004

**Keywords:** enteric glial cell, metabolic disorder, obesity, gut

## Abstract

Immune and nervous system sensing are two important ways of detecting inner and outer environmental changes. Immune cell activation in the gut can promote metabolic disorders. However, whether enteric nervous system sensing and activities are also important in metabolic syndromes is not clear. Enteric glial cells (EGCs) are thought to have sensing ability, but little is known about the potential connections between EGC and metabolic disorders. Consuming a modern Western-type high-fat low-fiber diet increases the risk of obesity. Here, we reported that dietary shift from a normal chow diet to a high-fat diet in wild-type (WT) C57BL/6 mice induced a transient emergence of glial fibrillary acidic protein (GFAP)-positive EGC network in the ileal lamina propria, accompanied by an increase of glial-derived neurotrophic factors production. Inhibition of EGC metabolic activity via gliotoxin fluorocitrate or glial-intrinsic deletion of myeloid differentiation factor 88 (*Myd88*) in mice blocked this dietary change-induced activity. Furthermore, we found a different role of MYD88 in glial cells versus adipocyte in diet-induced obesity. The glial *Myd88* knockout mice gained less body weight after HFD feeding compared to the littermate controls. In contrast, adipocyte deletion of *Myd88* in mice had no impact on the weight gain but had exacerbated glucose metabolic disorders. Pharmacological interventions of glial activities by fluorocitrate prevented body weight gain in a dietary type- and glial MYD88-independent manner. Collectively, our data reveal a previously unappreciated function of EGC in sensing a dietary shift-induced perturbation and glial activities as a whole may play roles in diet-induced obesity.

**New & Noteworthy:** It is known that obesity and its related metabolic syndrome can damage the neuronal system. However, whether the neuronal system also participates in the development of obesity is unclear. Diet is an important contributing factor to obesity. Our study reveals that consuming a high-fat diet can induce a transient enteric glial cell response via its intrinsic sensing molecule(s). Inhibiting overall glial cell activities may have an impact on the development of the metabolic syndrome.

## Introduction

Obesity is not simply a disease of abnormal body fat accumulation, it is often accompanied by a low-grade systemic chronic inflammation that can induce metabolic complications, including insulin resistance and type 2 diabetes (33). The gut is considered as an important contributor to metabolic diseases. It is the primary site for the absorption of food energy. Obesity predisposes to alter gut microbiota (27). The compositional and functional disturbances of the gut microbiota, i.e. dysbiosis, can promote obesity development. Transfer of gut flora from obese mice to germ-free mice increased obesity (27). In humans, it has been shown that intestinal transfer of fecal microbiota from lean donors can improve insulin sensitivity in recipients with metabolic syndrome (31). Obesity is also associated with dysfunctions of the intestinal barrier, which cause enhanced permeability and translocation of bacteria or bacterial products to the intestinal lamina propria and systemic circulation (1, 3). This influx of immune-stimulatory microbial ligands into the circulation has been suggested to drive the systemic chronic inflammation and metabolic syndrome accompanied with obesity (4). Furthermore, obesity predisposes to altered the intestinal immune system and the gut immune system has become a novel therapeutic target for metabolic diseases (33). The intestinal tract contains its own enteric nervous system (ENS), which includes a large number of neurons and even greater number of glial cells that control intestinal motor, sensory, absorptive, and secretory functions (30). However, whether there is a connection between ENS and obesity is still not clear.

Enteric glial cells (EGCs) are the major cellular component of the ENS and are found within the ganglia of the myenteric and submucosal plexus and in extraganglionic sites, such as the smooth muscle layers and the mucosa. EGCs are heterogenous and have shown phenotypic plasticity in the expression of some of the key feature surface molecules, such as glial fibrillary acidic protein (GFAP) (2). However, our current understanding of the physiological significance of EGC diversification remains unclear. EGCs play important roles in regulating intestinal motility and are even thought to be important in regulating epithelial barrier function, although the latter is still controversial (9, 19, 22). EGCs can engage functional crosstalk with all the different cell types present in the gut wall, including intestinal neurons, gut epithelium, enteroendocrine cells and immune cells, via direct physical contact or indirect paracrine secretions (28). EGCs express pattern recognition receptors, especially toll-like receptors (TLRs) (13, 20), and are able to perceive and integrate microbiota- and tissue-derived alarmin signals, in a manner somewhat dependent on the adaptor myeloid differentiation factor 88 (MYD88) (12). EGCs express neurotrophic factors, which can regulate intestinal homeostasis, inflammation, and defense against infection through crosstalk between EGC, immune cell, and the gut epithelium (12). Whether EGCs play any role in metabolic disease is not known. In this study, we set up experiments to explore the potential connections between EGC and metabolic disorders.

## Materials and Methods

### Animal studies

All our experimental procedures were approved by the Animal Research Committee of Xuzhou Medical University. The mice were housed at 20~22°C with a12h light-dark cycle and given free access to food and water. All animals were in the C57BL/6 (B6) background and maintained at our in-house animal facility free of specific pathogens. Only male mice were used in our experiments. The following animal *strains-Adiq-cre* (stock number 028020), *GFAP-cre* (stock number 012886), and *MyD88^fl/fl^* mice (stock number 008888) were purchased from the Jackson Laboratory (Bar Harbor, ME, USA). WT B6 mice were from Beijing Vital River Laboratory Animal Technology Co., Ltd (Beijing, China). High-fat diet (HFD), 60% of calories from fat (D12492; Research Diets), was purchased from Beijing Keao Xieli Feed Co., Ltd (Beijing, China).

### Fluorocitrate preparation

Fluorocitrate was prepared following a previously described method (21). Briefly, DL-fluorocitric acid barium salt (Sigma-Aldrich Co., St. Louis, MO, USA) was dissolved in 0.1 M HCl and the Ba^2+^ was precipitated by the addition of 0.1 M Na_2_SO_4_. This solution was clarified by centrifugation at 800 g for 10 min and the supernatant removed and diluted in saline solution (NaCl, 0.9%) to a final concentration of 100 μM (adjusted pH 7.0 by addition of 1 N NaOH). For FC treatment, 20 μmol/kg FC was administered intraperitoneally twice daily. For treatment control, some groups of mice were injected with PBS.

### Quantitative RT-PCR

Total RNA was extracted from tissues that homogenized in Trizol reagent (Thermo Fisher Scientific, Waltham, MA, USA). One microgram of purified RNA from each sample was used for reverse transcription reaction to generate cDNA with a high-capacity cDNA reverse transcription kit (Takara, Dalian, China) and the resulting cDNA was then used for qPCR on a real-time PCR detection system (Bio-Rad, Hercules, CA, USA). The relative mRNA expression level was determined with the 2^-ΔΔCt^ method, using *β-actin* as the internal reference control. Primer sequences were as the following: *artn*(*F*: 5’-GCACACTAGACCCATGTGTTGC-3’, R:5’-CTCCCAGAGGAGTTCTCTTTAGC-3’); *gdnf*(F:5’-GGGGTGCGTTTTAACTGCC-3’, R:5’-GTTTAGCGGAATGCTTTCTTAG-3’); *s100β*(F:5’-AAGATGGGGATGGGGAGTG-3’, R:5’-CAAAGCAAACCAAGCTTCC-3’); *β-actin*(F:5’-TATTGGCAACGAGCGGTTC-3’, R:5’-ATGCCACAGGATTCCATACCC-3’).

### Immunoblot

Tissue homogenates were collected in RIPA buffer containing 50 mm Tris (pH 7.4), 150 mM NaCl, 1% Triton X-100,1% sodium deoxycholate, 0.1% SDS with freshly added protease inhibitors and phosphatase inhibitors. They were then clarified by centrifugation at 12,000rpm at 4°C for 15 min. The protein concentration in the supernatant was determined by bicinchoninic acid (BCA) method. Proteins were separated by SDS-PAGE electrophoresis and transferred to PVDF membrane. Target proteins were reacted with primary antibodies after blocking PVDF membranes with 5% skimmed milk for 1 hr. After incubating with the secondary antibody (horseradish peroxidase-conjugated anti-rabbit IgG), immunoreactive bands were detected with enhanced chemiluminescence system (Pierce, Rockford, IL, USA). The rabbit primary Abs against the following proteins were used: Artemin and S100β (Abcam, Cambridge, United Kingdom), and β-actin (Abclonal, Wuhan, China). The band intensities on the blot were quantified with ImageJ software (National Institutes of Health, Bethesda, MD, USA).

### Immunofluorescence

For immunostaining, ileal segments were fixed in 4% paraformaldehyde at room temperature for 48 h and embedded in paraffin. Serial sections (4 μm) were obtained, deparaffinized in xylene, and rehydrated in graded alcohol solutions. Antigen retrieval was performed with EDTA antigen retrieval buffer (Beyotime, Shanghai, China). The sections were then blocked in 3% BSA for 30 min. Afterward the sections were incubated with 1:100 diluted mouse polyclonal anti-GFAP (Cusabio, Houston, TX, USA) and 1:500 diluted rabbit polyclonal anti-HuC/D (Abcam, Cambridge, United Kingdom) overnight at 4°C· followed by incubation with Cy3-labeled goat anti-mouse IgG (Beyotime, Shanghai, China) and Alexa Fluor 488-labeled goat anti-rabbit IgG (Beyotime, Shanghai, China) for 50 min at RT. Nuclei were counterstained with 4’, 6-diamidino-2-phenylindole, dihydrochloride (DAPI) (Beyotime, Shanghai, China). The images were collected under a fluorescence microscope.

### Isolation of enteric glia cells from the myenteric plexus

EGCs were isolated according to a previously published protocol (24) with minor modification. Briefly, mice were euthanized and the intestine from the duodenum to the ileum was unraveled. The intestinal segments were washed by cold phosphate buffer (PBS), and cut into 3 cm of small segments. The longitudinal muscle/myenteric plexus (LMMP) of the intestine was harvested by creating a gap in the longitudinal muscle and using a moist cotton swab to rub horizontally. LMMP was placed into a digestion solution (collagenase type II 13mg, BSA 3mg, dissolved in 10ml PBS), and digested at 37°C for 60 min. After centrifugation, the cell pellet was digested by 0.05% trypsin solution at 37°C for 7 min. The mixture was filtered through a Nitex nylon mesh and centrifuged at 356 g for 8 min. The precipitate was resuspended in enteric neuron media (neurobasal A media with B27, 2 mM L-glutamine, 1% FBS, Glial-derived neurotrophic factor (GDNF) 10 ng/ml, and an antibiotic/antimycotic cocktails, all from Thermo Fisher Scientific, Waltham, MA, USA). The cells were cultured for 7 days at 37°C, 5% CO_2_. To determine the expression level of MYD88 in the EGCs, we used an anti-MYD88 antibody from Abcam (Cambridge, UK).

### Glucose tolerance test (GTT)

For the GTT assay, the mice were fasted overnight and then injected intraperitoneally with glucose solution at 2 g/kg. Blood was collected at 0, 30, 60, 90 and 120 min accordingly.

### Statistical analysis

Data were analyzed with GraphPad Prism 6.0 (GraphPad Software, La Jolla, CA, USA) and are presented as means ± SE. Statistical significance was determined using the unpaired two-tailed Student’s t-test for two datasets and one-way or two-way analysis of variance (ANOVA) followed by Bonferroni post hoc tests for multiple groups. *P* < 0.05 was considered statistically significant.

## Results

### Dietary change initiates a short-term enteric glial cell response

Modern industrialization makes high-fat, low-fiber Western-type diet become popular, and this type of diet increases the risk of fat mass accumulation and promotes obesity (26). How the host responds to such type of diet remains incompletely understood. To explore whether EGC can sense a dietary change, 6~8-week-old wild-type (WT) B6 male mice that were previously fed with a normal chow diet (ND) were switched to a high-fat diet (HFD) on day 0, the potentials of glial cell response were monitored thereafter by examining glial cell marker gene’s mRNA and protein expression in tissue segments from the proximal duodenum to the distal colon by qRT-PCR and immunoblot, respectively. As shown in Fig. 1A, the calcium-binding protein *S100β*, glial cell-derived neurotrophic factor (*Gdnf*), and GDNF family member artemin (*Artn*) mRNA expression levels were all significantly up-regulated specifically in the ileum 4~6 days after the dietary shift. Ileal S100β and ARTN protein expression levels were also elevated at day 3 after HFD feeding (Fig 1B and 1C). In order to confirm that the increased expression of S100β and ARTN were indeed derived from EGC, we intraperitoneally injected fluorocitrate (FC), a selective inhibitor of EGC activities (6, 18), to the mice twice daily for 3 consecutive days starting from the beginning of HFD feeding. The increment of ileal S100β and ARTN expressions by HFD feeding was blocked after the FC treatment (Fig.1B and 1C). These results indicate that the dietary shift induces an EGC response in the ileum.

**Figure1.**
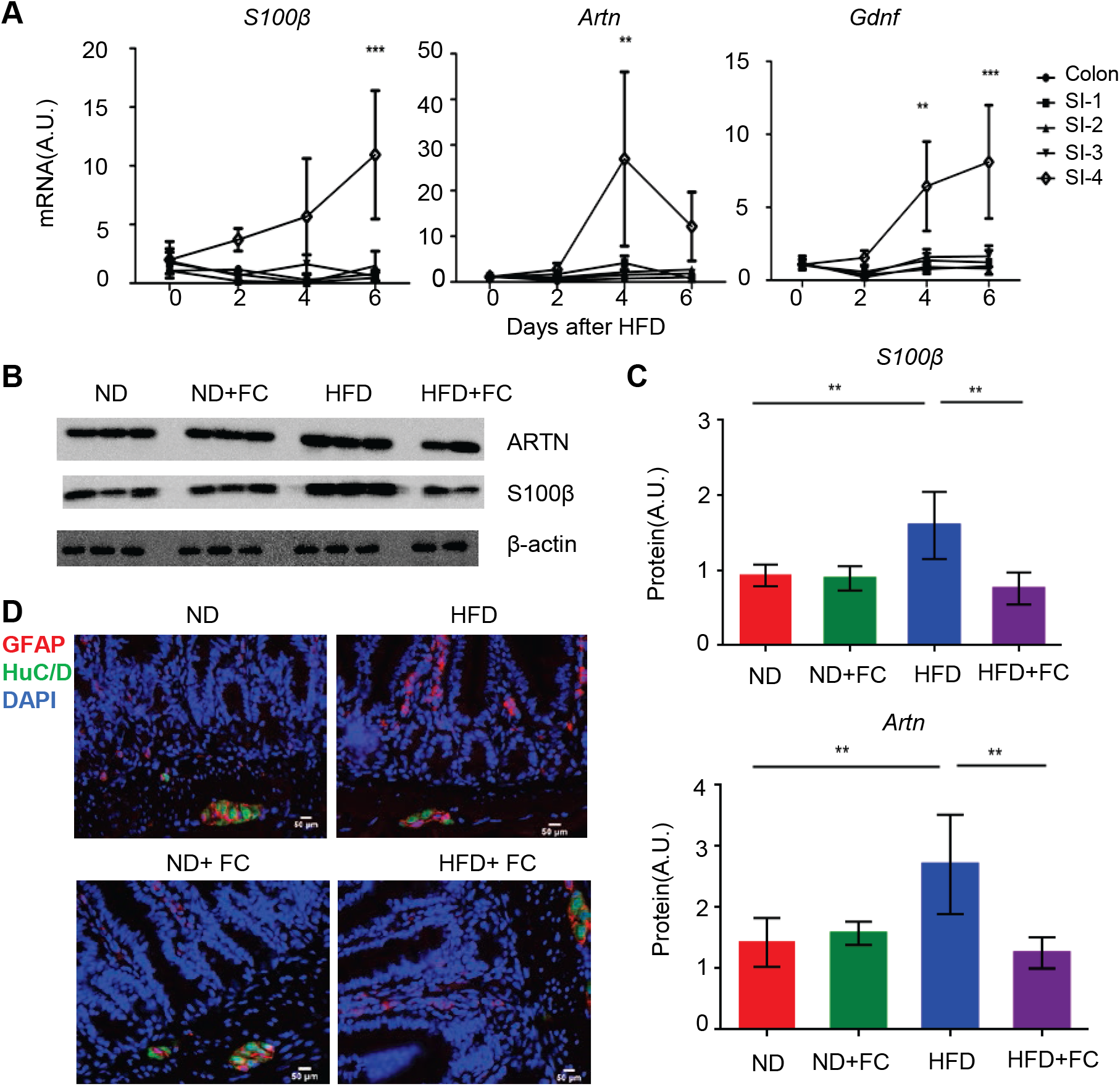
Dietary change from a normal chow diet (ND) to a High-fat diet (HFD) induced an enteric glial cell response. 6-8-week-old WT B6 male mice initially fed with ND were switched to HFD on day 0. **(A)** Time course of the relative mRNA expression of *s100β, artn*, and *gdnf* in the indicated intestinal tissue segments after HFD feeding (n=3-5 in each time point). The small intestine (SI) was manually divided into 4 roughly equal parts, the segments were labeled sequentially from 1 to 4, starting from the proximal duodenum (SI-1) to the distal ileum (SI-4).\*\**P* < 0.01; \*\*\**P* < 0.001 by two-way ANOVA plus Bonferroni post hoc tests. **(B)** Immunoblot analysis of the ileal S100β and ARTN expression at day 3 after HFD consumption. Fluorocitrate (FC) was administered twice daily for a total of three days starting from day 0. Each lane represents a sample from one individual mouse. **(C)** Densitometry quantification for (B). ** *P* < 0.01 by one-way ANOVA plus Bonferroni post hoc tests. **(D)** Immunofluorescence for glial cell marker GFAP and pan-neuronal marker HuC/D in the ileal tissue sections derived from the wild-type B6 mice that were either on ND or switched to HFD for 3 days in the presence or absence of FC treatment. Scale bar = 50 μm.

EGCs are heterogeneous populations and have shown phenotypic plasticity (2). To explore which populations of EGC respond to the dietary change, we performed immunohistology analysis. There was an emergence of glial fibrillary acidic protein (GFAP)-positive cell network in the lamina propria along with the ileal villus-crypt units 3 days post HFD, which was usually not seen under ND condition (Fig. 1D). The emergence of these cells was effectively blocked by the FC treatment (Fig. 1D). In contrast, the myenteric glial cells that located within the myenteric ganglia and the extraganglionic muscular layers exhibited little change of their GFAP expression, and their GFAP expression appeared to be insensitive to the FC treatment (Fig. 1D). The data collectively suggest a mucosal EGC alteration after the dietary shift.

The dietary change-induced EGC response is transient. The upregulated S100β and ARTN expressions in the ileum were not sustained. 12 weeks after HFD feeding, S100β and ARTN expression levels became similar to that of the ND control (Fig. 2A and 2B). GFAP expression in the lamina propria also returned to the background level (Fig. 2C). Interestingly, the GFAP and HuC/D expression levels within the myenteric ganglia were very stable and showed minimal changes after long-term HFD diet feeding, and they were not influenced even by 12 weeks of consecutive FC treatment (Fig. 2C).

**Figure 2.**
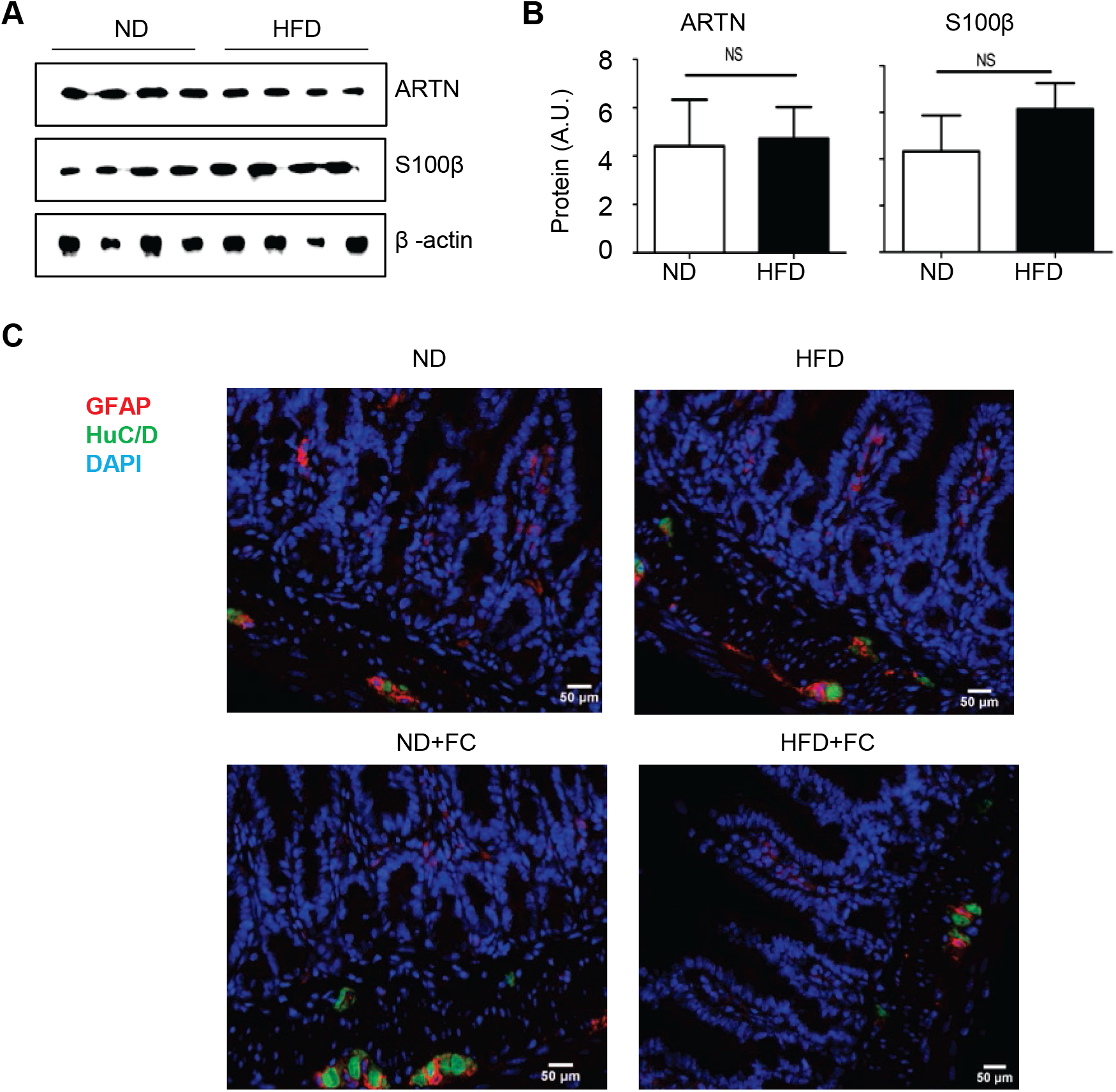
The dietary shift-induced mucosal EGC response was diminished after long-term HFD feeding. WT B6 mice initially fed with ND were either continuous to fed with ND or switched to HFD on day 0. **(A)** Immunoblot analysis of ileal S100β and ARTN expression in week 12 post HFD feeding. Each lane represents a sample from one individual mouse. **(B)** Densitometry quantification for (A). One representative experiment of at least 3 independent experiments is shown. Data are presented as mean ± SE (total accumulated n=9-15 in each group). Ns, no statistical significance by two-tailed Student’s t-test. **(C)** WT B6 mice were fed with ND or HFD for 12 weeks in the presence or absence of 12-week daily FC treatment as indicated. Representative immunofluorescence image for glial cell marker GFAP and pan-neuronal marker HuC/D in the ileal tissue sections derived from the indicated mice were shown. Scale bar = 50 μm.

### The dietary change-induced EGC response requires MYD88

It has been reported that EGCs can sense commensal products and/or alarmins to produce neurotrophic factors via the cell-intrinsic MYD88-dependent pathway (12). To interrogate whether the dietary shift-induced glial cell response is MYD88-dependent, we deleted *Myd88* in GFAP expressing glial cells by crossbreeding *Gfap*-Cre to *Myd88^fl/fl^* mice (11, 25). Glial-intrinsic deletion of *Myd88* was confirmed by western blot analysis of the MYD88 expression in ex-vivo isolated myenteric glial cells from the corresponding mice (Fig. 3A). Although GFAP-cre marks only a relatively small subset of EGCs (15), remarkably, the dietary shift-induced EGC response was significantly attenuated in the *Gfap-Cre/Myd88^fl/fl^* mice (Fig. 3B and 3C). Furthermore, FC treatment showed a minimal impact in further reducing the EGC response in these mice (Fig. 3D-3F). The data suggest that the glial cell-intrinsic MYD88 sensing pathway is required for the dietary change-induced EGC response.

**Figure 3.**
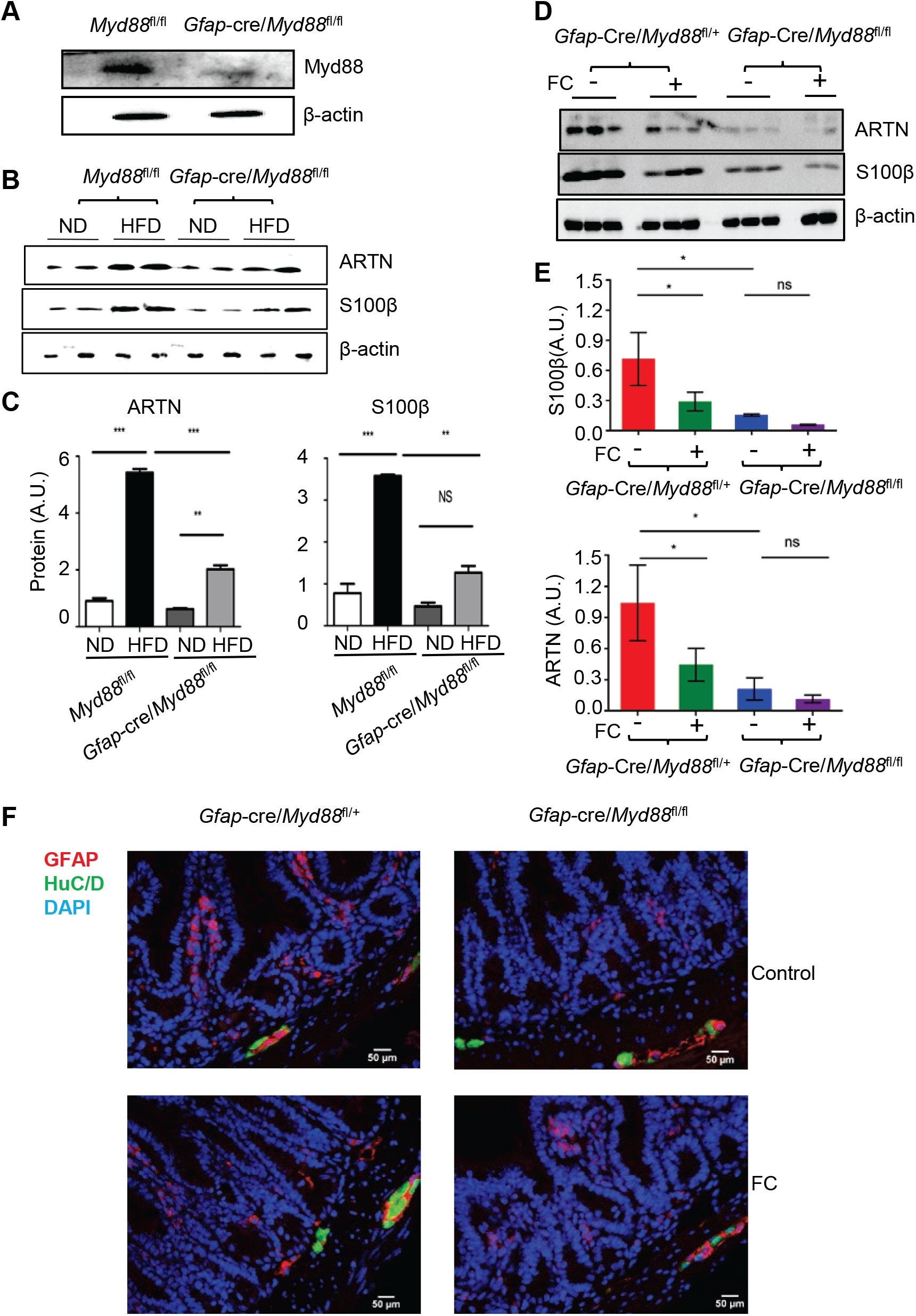
EGC intrinsic MYD88 pathway participated in the dietary change-induced EGC response. **(A)** Immunoblot analysis of MYD88 expression in ex-vivo purified EGCs collected from the indicated mice. **(B-F)** 6~8-week-old *Gfap*-cre/*Myd88^fl/fl^* male mice and their littermate controls initially fed with ND were switched to HFD on day 0, and in the meantime, they were treated with vehicle or FC twice daily for a total of 3 days. (B and C) immunoblot analysis of the ileal S100β and ARTN expression on day 3 post HFD (B), and their densitometry quantification (C). \*\**P* < 0.01; \*\*\**P* < 0.001; ns, no statistical significance by one-way ANOVA with Bonferroni post hoc tests. (D and E) immunoblot analysis of the effect of FC treatment on the ileal S100β and ARTN expression at day 3 post-HFD in the indicated mice (D). (E) densitometry quantification for (D). \**P* < 0.05; ns, no statistical significance by one-way ANOVA with Bonferroni post hoc tests. (F) Immunofluorescence for glial cell marker GFAP and pan-neuronal marker HuC/D in the ileal tissue sections derived from the indicated mice. Scale bar = 50 μm.

### Glial-intrinsic Myd88 sensing pathway participates in regulating HFD-induced body fat accumulation

To interrogate the physiological importance of MYD88-dependent glial cell signals on the HFD-induced obesity, 6~8-week-old *Gfap-Cre/Myd88^fl/fl^* and their littermate *Gfap-Cre/Myd88^fl/+^* male mice previously fed on ND were switched to HFD on day 0 and were maintained on HFD for 12 weeks. The two groups of mice had similar body weight initially. After HFD, *Gfap-Cre/ Myd88^fl/fl^* mice gained less weight on a weekly basis relative to the littermate controls (Fig. 4A), although their food intake was similar (Fig. 4B). Twelve weeks after HFD, *Gfap-Cre/Myd88^fl/fl^* mice accumulated less body fat (Fig. 4C). Interestingly, although at this time *Gfap-Cre/Myd88^fl/fl^* mice were leaner, their glucose tolerance capacity had no improvement (Fig. 4D and 4E). These results suggest that the glial cell MYD88-dependent sensing pathway participates in HFD-induced obesity. However, whether this is related to the diet-induced mucosal EGC response remains to be determined.

**Figure 4.**
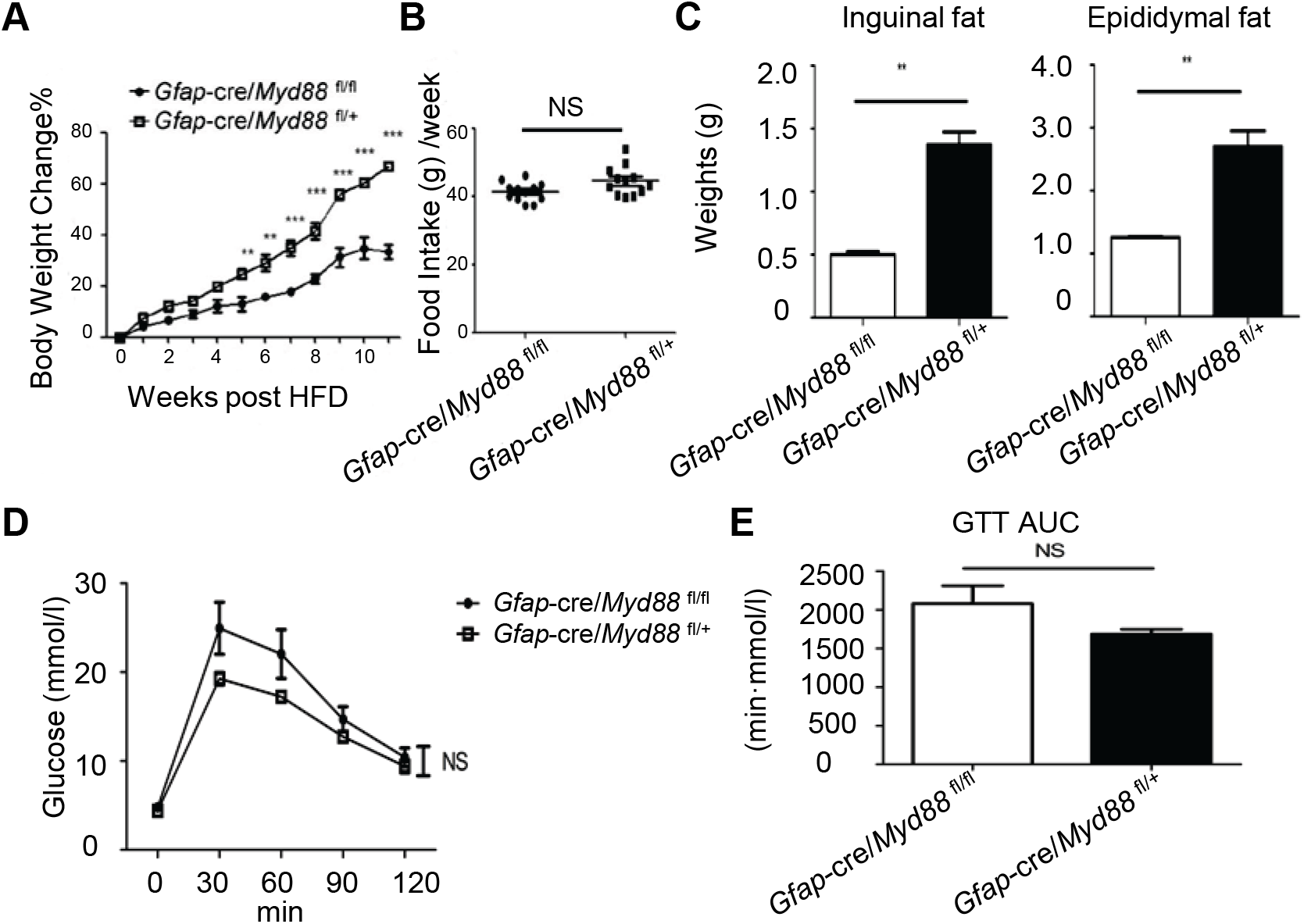
Glial-intrinsic deletion of *Myd88* reduced diet-induced fat mass accumulation. 6~8-week-old *Gfap*-cre/*Myd88^fl/fl^* male mice and its littermate controls were fed with HFD for 12 weeks. **(A)** Weekly body weight changes. \*\**P* < 0.01; \*\*\**P* < 0.001 by two-way ANOVA plus Bonferroni post hoc tests. **(B)** Weekly food intake. NS, no statistical significance by T-test. **(C)** Fat-pad mass at 12 weeks post HFD. \*\**P* < 0.01 by T-test. **(D and E)** After 12 weeks of HFD feeding, mice were fasted overnight and administered glucose intraperitoneally. Blood glucose levels were measured at the indicated time point (D) and the area under curve (AUC) was calculated accordingly (E). The statistics were evaluated by two-way ANOVA (D) and T-test (E), respectively. One representative experiment of 2 independent experiments is shown. Data are presented as mean ± SE (n=6-8 in each group).

### Adipocyte-intrinsic Myd88 sensing pathway modulates HFD-induced glucose metabolic disorders

It is considered that high-fat diet-induced systemic endotoxemia is an important driver in promoting obesity and its related complications (3, 7). Although our data so far imply a potential link between glial sensing and diet-induced obesity, whether MYD88-dependent sensing of commensal products or alarmins in adipocyte directly impacts the development of obesity is not known. To this end, we deleted Myd88 in adiponectin (Adiq) expressing adipocyte by crossbreeding *Adiq-Cre* to *Myd88^fl/fl^* mice. Adipocyte-intrinsic deletion of Myd88 did not influence the mice on weight gain either on HFD-or ND-fed condition (Fig. 5A) and did not show an impact on fat mass accumulation after 3 months of HFD consumption (Fig. 5B). The amount of weekly food intake also did not change (Fig. 5C). Interestingly, although there was no bodyweight difference, *Adiq-Cre/Myd88^fl/fl^* mice displayed worse glucose intolerance than the *Myd88^fl/fl^* littermate control (Fig 5D and 5E). These results suggest that, in contrast to the role of glial cell-derived MYD88 pathway in promoting fat mass accumulation, adipocyte intrinsic MYD88 pathway may instead function as a limiting factor in diet-induced metabolic abnormalities.

**Figure 5.**
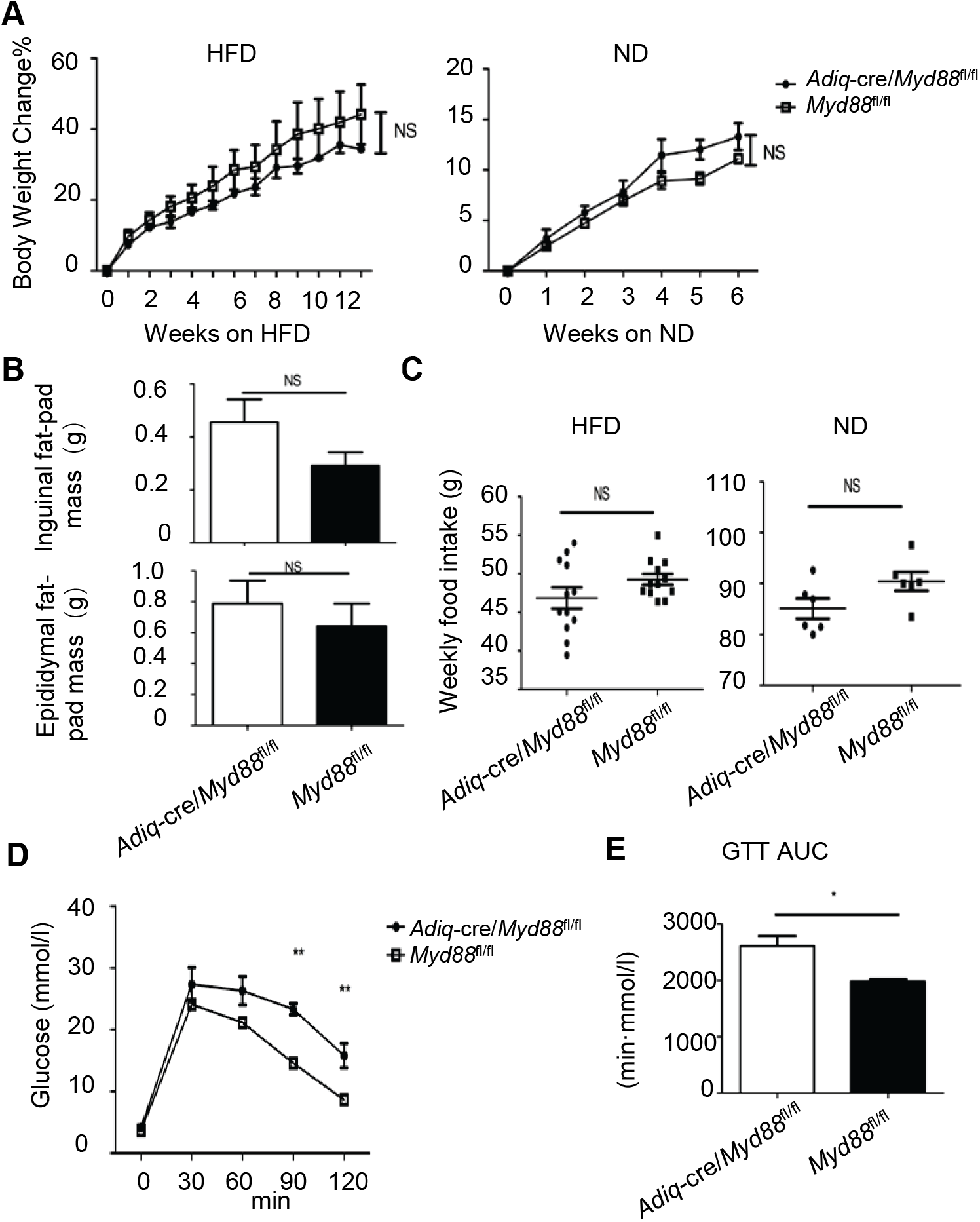
Adipocyte-intrinsic deletion of Myd88 promoted diet-induced glucose metabolic abnormalities. 6~8-week-old *Adiq*-cre/*Myd88^fl/fl^* male mice and its littermate controls previously fed on ND were switched to HFD. As controls, we used mice that were continuously fed with ND. **(A)** The body weight changes with time. NS, no statistical significance by two-way ANOVA. **(B)** Fat-pad mass at 12 weeks post-HFD. NS, no statistical significance by T-test. **(C)** Weekly food intake. NS, no statistical significance by T-test. **(D and E)** After 12 weeks of HFD feeding, the mice were fasted overnight and then administered glucose intraperitoneally. Blood glucose levels were measured at the indicated time point (D) and the area under curve (AUC) was calculated accordingly (E). \**P* <0.05; \*\**P* < 0.01 by two-way ANOVA (D) and T-test (E), respectively. One representative experiment of 2 independent experiments is shown. Data are presented as mean ± SE (n=6-8 in each group).

### Pharmacological intervention of glial activities by fluorocitrate prevented body weight gain in a dietary type- and glial MYD88-independent manner

To further explore whether interruption of glial cell function pharmacologically can impact diet-induced obesity, groups of WT B6 mice previously fed on ND were either continue to be fed with ND or switched to HFD on day 0, in the meantime treated with vehicle or FC intraperitoneally twice daily for a week and then the treatment was discontinued. When FC treatment was applied, the mice immediately stopped gaining weight on both ND- and HFD-fed conditions (Fig. 6A). However, once FC treatment was terminated, the mice started to regain weight (Fig. 6A), implying a transitory and reversible nature of the FC treatment. There was no post-FC anorexia, indicated by relative food intake measurement (Fig. 6B). 4 weeks after FC termination no alleviation of HFD-induced glucose intolerance was observed (Fig. 6C and 6D), but the epidydimal fat mass accumulation induced by the HFD was reduced (Fig. 6E and 6F).

**Figure 6.**
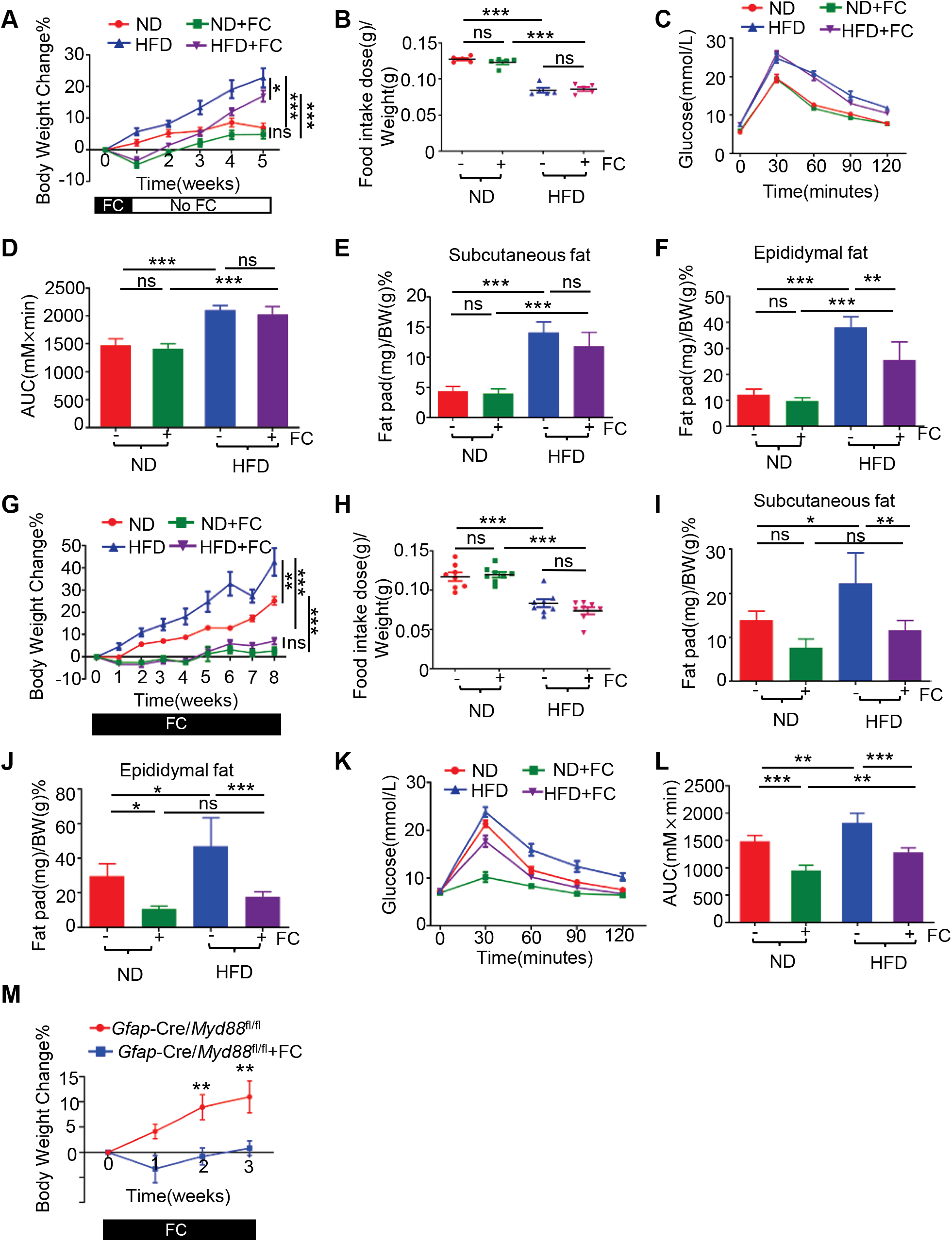
Pharmacological intervention of glial activities by fluorocitrate prevented body weight gain. **(A-L)** Groups of wild-type B6 mice previously fed on ND were either continue to be fed with ND or switched to HFD on day 0. (A-F) Mice were treated with FC or vehicle intraperitoneally for 1 week starting from day 0 and then the treatment was discontinued. (A) Weekly body weight changes. The statistics were evaluated by two-way ANOVA with Bonferroni post hoc tests. (B) Relative food intake per week. The statistics were evaluated by one-way ANOVA with Bonferroni post hoc tests. (C) Glucose tolerance test. (D) The area under curve (AUC) for (C). The statistics were evaluated by one-way ANOVA with Bonferroni post hoc analysis. (E, F) The inguinal (E) and epididymal (F) fat-pad mass in the indicated mice 4 weeks after FC termination. The statistics were evaluated by one-way ANOVA with Bonferroni post hoc analysis. (G-L) Mice were treated with FC or vehicle consecutively for a total of 8 weeks starting from day 0. Similar statistical methods were used as in (A-F), respectively. (G) Weekly body weight changes. (H) Relative food intake per week. (I, J) The inguinal (I) and epididymal (J) fat-pad mass 8 weeks after FC treatment. (K) Glucose tolerance test. (L) The area under curve (AUC) for (K). **(M)** 6~8-week-old *Gfap-cre/Myd88^fl/fl^* male mice initially fed on ND were switched to HFD on day 0 and in the meantime were treated with vehicle or FC twice daily for a total of 3 weeks. The weekly bodyweight changes were shown. For all the graph, \**P* < 0.05; ** *P* < 0.01; \*\*\**P* < 0.001; ns, no statistical significance. (n=5 in each group).

When we extended the FC treatment to 8 weeks without interruption, we found it could completely prevent body weight gain on both ND- and HFD-fed condition during the whole treatment period (Fig. 6G). Switching ND to HFD could reduce the relative food intake, but FC treatment per se did not have an impact (Fig. 6H). Long-term FC treatment prevented body fat accumulation (Fig. 6I and 6J) and significantly improved glucose tolerance 8 weeks post HFD (Fig. 6K and 6L). These results together suggest of potentials of targeting glial activities for metabolic disorders.

Finally, the FC treatment was still effective in inhibiting weight gain in the *Gfap-Cre/Myd88^fl/fl^* mice (Fig. 6M). Thus, the effect of FC on weight gain was glial MYD88-independent.

## Discussion

Recent research progress has pinpointed the gut as a critical contributor and an important therapeutic target for metabolic disorders (33). Although the enteric immune system has been recognized as an important contributor to metabolic disorders, the enteric nervous system is present in the gut in large numbers, but very little is known about their roles in metabolic diseases. In this study, we reported that dietary shift from a normal chow diet to an energy-dense high-fat low fiber diet in mice induced a transient glial cell response in the ileal lamina propria. Either blocking EGC metabolic activity via fluorocitrate or deletion of Myd88 in GFAP-expressing cells inhibited this response. Our data support a notion of ENS, especially EGCs, as a critical gatekeeper that constantly monitors and responds to environmental challenges from the gut ecosystem, to modulate intestinal homeostasis.

What exactly EGCs sense during the dietary shift remains to be determined. A previous study by Kabouridis et. al. indicated that the mucosal glial cell networks are somewhat dependent on the gut microbiota (13). It is possible that EGC may sense newly derived microbial or dietary ingredients or their metabolites post dietary change.

Which type of glial cells sense the dietary change needs to be further studied. Although the dietary shift from a normal chow to a high-fat diet induces mucosal GFAP^+^ EGC accumulation, these cells might be originated from progenitor cells located in the ganglionic plexus as a previous study suggested (13). It is possible that the progenitor cells can sense the dietary change and differentiate and migrate to the villi. Another possibility is that the mucosal EGCs themselves sense and respond in situ. Given the fact that the mucosal EGCs locate in close proximity to the intestinal lumen, they are better suited to respond to changes happening in the lumen.

What the function of EGC sensing is during the dietary change remains unclear. Either the use of GFAP-cre mice or the administration of fluorocitrate have technical limitations and are not specific for targeting only the mucosal GFAP-positive EGCs. More advanced tools are needed to further address this question. Regardless of the limitations, our data imply that targeting overall glial cell activities (locally and/or systemically) can impact the host’s metabolism.

One possible scenario of mucosal EGC sensing might be regulating gut microbiota. EGC can release neurotrophic factors, such as GDNF and ARTN. These neurotrophic factors can activate innate lymphoid-like cell type 3 (ILC3) to produce interleukin-22 (IL-22) (12). IL-22 can stimulate the gut epithelium to produce anti-microbial peptides and these anti-microbial peptides can promote dysbiosis in the gut (5). Dysbiotic microbiota attains an increased capacity to harvest energy from the diet (27), which in short-term can be beneficial for the host to absorb dietary nutrients. However long-term dysbiosis with HFD will lead to obesity.

Much of the innate immune system sensing is governed by pattern recognition receptors (PRR), such as transmembrane surface or endosome toll-like receptors (TLRs) and cytosolic nucleotide-binding oligomerization domain (NOD)-like receptors (NLRs) that bind to microbe-associated molecular patterns (MAMPs) expressed by normal microbiota or pathogens. Like the innate immune cells, EGCs express PRRs, especially TLRs (13, 20). Thus, the intestinal immune system and EGCs share some common regulatory mechanisms, and they probably work as a whole to face gut environmental changes. This is consistent with the concept that neuroimmune cells can form a functional unit and work together in regulating intestinal development, homeostasis, and disease (29). EGCs can also interact with various other cells, such as neurons, enterocytes, enteroendocrine cells, muscle cells, and endothelial cells (8), how might those interactions influence obesity is still an open question. A most recent finding implies that some microbially responsive neurons in the distal ileum and proximal colon autonomously regulate blood glucose, however, they do not express pathogen recognition receptors, including TLRs (17). It is possible that the ileal mucosal EGCs respond to certain microbial-related signals and relay the information to the neuron to modulate glucose metabolism.

The role of MYD88 in obesity is currently not understood very well. There are conflicting reports in the literature (10, 14, 23, 32). We found a different role of MYD88 in glial cell versus adipocyte in diet-induced obesity. The deletion of glial MYD88 in mice primarily reduced body fat mass accumulation, but not diet-induced glucose intolerance. In contrast, the deletion of MYD88 in adipocyte promoted glucose intolerance, but not body fat mass accumulation. It implies a division of labor among the sensing cells to form an intricated regulatory network in maintaining the host’s metabolic homeostasis. Although obesity and metabolic disorders are often seen together, they are different processes and are regulated differently. Indeed, people with obesity can be metabolic healthy (16). Understanding the common and differential regulatory processes that control obesity and metabolic syndromes are critical for better-targeted strategies to manipulate these disorders.

In summary, our findings reveal a previously unappreciated function of EGC in sensing a dietary shift-induced perturbation and glial activities as a whole may play roles in diet-induced obesity.

## Disclosures

The authors declare that they have no competing interests

## Grants

Project support was provided in part by the National Natural Science Foundation of China (81770853 to YgW, 82002157 to ZL), the Priority Academic Program Development of Jiangsu Higher Education Institutions (PAPD) in the year of 2014 (Grant No. KYLX14-1448), Natural Science Foundation of the Higher Education Institutions of Jiangsu Province, China (Grant No. 16KJB310016), the Starting Foundation for Talents of Xuzhou Medical University (Grant No. D2016029).

---

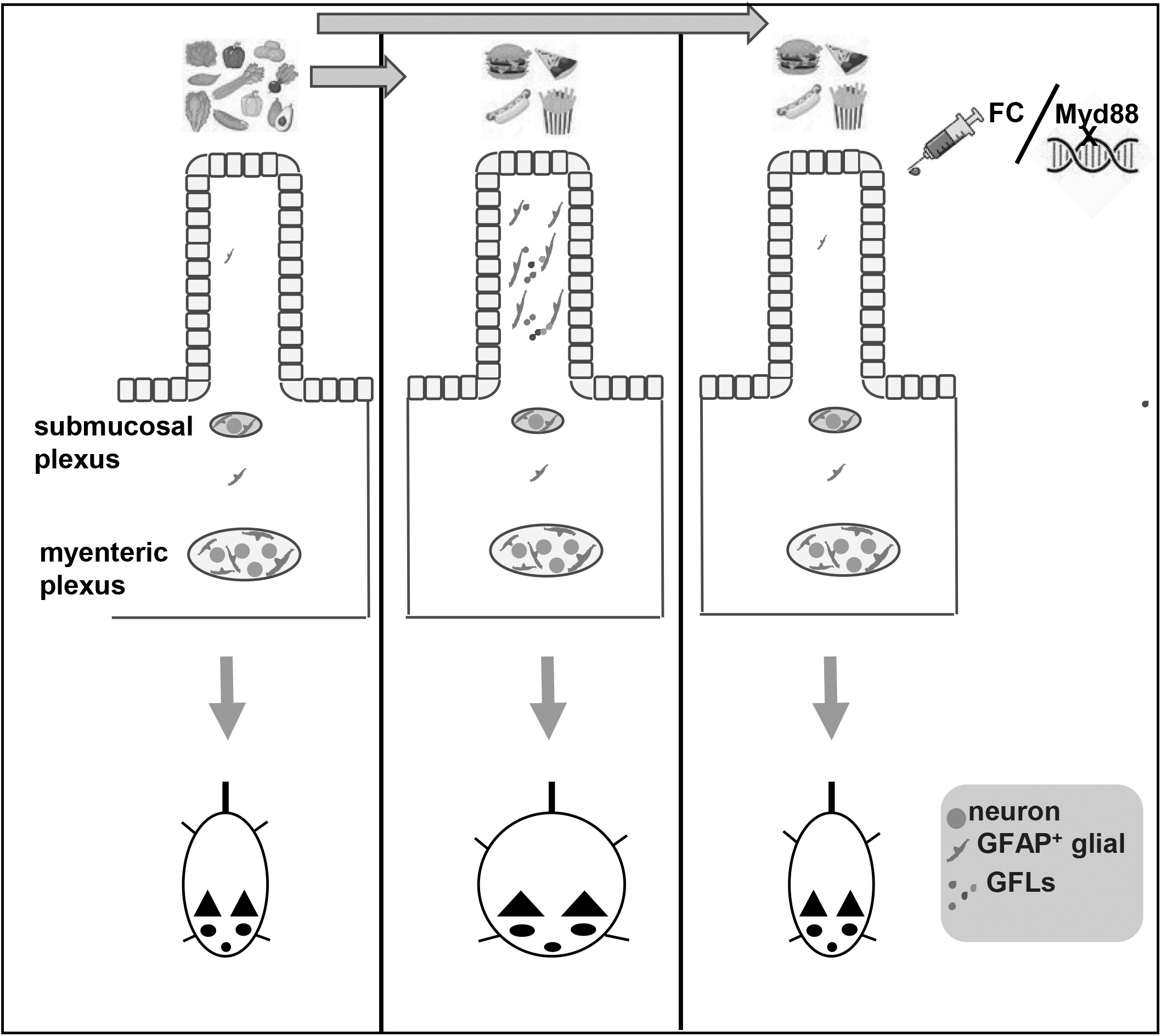

## Notes

### Competing Interest Statement

The authors have declared no competing interest.

